# A calcium-based plasticity model predicts long-term potentiation and depression in the neocortex

**DOI:** 10.1101/2020.04.19.043117

**Authors:** Giuseppe Chindemi, Marwan Abdellah, Oren Amsalem, Ruth Benavides-Piccione, Vincent Delattre, Michael Doron, Andras Ecker, James King, Pramod Kumbhar, Caitlin Monney, Rodrigo Perin, Christian Rössert, Werner Van Geit, Javier DeFelipe, Michael Graupner, Idan Segev, Henry Markram, Eilif Muller

## Abstract

Long-term potentiation (LTP) and long-term depression (LTD) of pyramidal cell connections are among the key mechanisms underlying learning and memory in the brain. Despite their important role, only a few of these connections have been characterized in terms of LTP/LTD dynamics, such as the one between layer 5 thick-tufted pyramidal cells (L5-TTPCs). Comparing the available evidence on different pyramidal connection types reveals a large variability of experimental outcomes, possibly indicating the presence of connection-type-specific mechanisms. Here, we show that a calcium-based plasticity rule regulating L5-TTPC synapses holds also for several other pyramidal-to-pyramidal connections in a digital model of neocortical tissue. In particular, we show that synaptic physiology, cell morphology and innervation patterns jointly determine LTP/LTD dynamics without requiring a different model or parameter set for each connection type. We therefore propose that a similar set of plasticity mechanisms is shared by seemingly very different neocortical connections and that only a small number of targeted experiments is required for generating a complete map of synaptic plasticity dynamics in the neocortex.

## Introduction

Neurons in the brain interact with each other via a complex network of synaptic connections, constantly adapting to external stimuli and internal dynamics. This remarkable capability, commonly referred to as “synaptic plasticity”, is thought to be the foundation of learning and memory in the brain (Bliss and Collingridge, 1993). Long-term potentiation (LTP) and long-term depression (LTD) of synaptic responses are the most basic forms of persistent connectivity adaptation (Bliss and Lømo, 1973; Lynch et al., 1977; Dunwiddie and Lynch, 1978).

LTP/LTD have been extensively studied for the connection between layer 5 thick-tufted pyramidal cells (L5-TTPCs) in the neocortex (Markram et al., 1997b; Sjöström et al., 2001, 2003, 2007) and a few other neocortical connection types (Egger et al., 1999; Sjöström and Häusser, 2006; Rodríguez-Moreno and Paulsen, 2008). These studies show that the outcome of plasticity experiments is highly diverse, depending on the pre- and post-synaptic neuron types involved. For this reason, it could be assumed that different plasticity rules exist between different connection types. However, due to the complexity and limitations of experimental procedures, most connection types remain unexplored with respect to plasticity. How then are we to get a comprehensive view of the plasticity dynamics governing the learning in neocortical circuits? Furthermore, the bulk of LTP/LTD data is made by somatic recordings of postsynaptic potentials (PSPs). As neocortical connections are generally mediated by multiple synapses (Markram et al., 1997a; Feldmeyer et al., 1999, 2006), somatic recordings only offer a superimposed view of all synaptic changes occurring at a connection. That is, even for the connection types that were experimentally characterized, we do not know what is happening at the single synapse level: are all synapses in a connection changing in a similar manner? Under what circumstances does the behavior of two synapses in the same connection diverge? How much does the current state of an individual synapse affect its capability to change? For all these reasons, studying learning and memory in terms of punctual LTP/LTD events is still very challenging.

Several studies have shown that synapse location on the dendrites plays an important role in determining the direction and magnitude of synaptic changes (Froemke et al., 2005; Sjöström and Häusser, 2006). As the distribution of dendritic locations of synapses is highly specific to pre- and post-synaptic neuron type, it is possible that this specificity could account for some of the specificity and diversity in plasticity outcomes, through the established dependence of plasticity on dendritic location. In this work, we investigated to what extent variability in innervation patterns alone combined with a single mechanism for LTP/LTD can account for the observed diversity in plasticity outcomes between the various neuron types. We propose a calcium-based model of LTP/LTD at the single synapse level, accounting for the heterogeneity of synaptic physiology. We show that this synaptic rule parameterized to model the synapses on L5-TTPCs can be transplanted as-is to describe synaptic plasticity data for other pyramidal– to–pyramidal synapses in the somatosensory cortex. This observation suggests that a similar set of plasticity mechanisms is shared among excitatory neocortical synapses, and that only a small number of targeted experiments could be sufficient to generate a complete map of synaptic plasticity dynamics in the neocortex.

## Methods

As previously mentioned, the behavior of individual synapses during LTP/LTD is not directly observable in most *in vitro* experiments, where only somatic PSPs can be measured (i.e. Markram et al. (1997b); Sjöström et al. (2001)). For this reason, how LTP/LTD mechanisms operate at these synapses is still a working hypothesis and an extensive description of those is beyond the scope of this work (for a review see Malenka and Bear (2004)). However, the general traits are now consolidated in literature and here briefly illustrated (Figure 1A). It is widely accepted that postsynaptic calcium, entering dendritic spines via N-methyl-D-aspartate receptors (NMDARs) and voltage-dependent calcium channels (VDCCs), is the key signaling molecule for LTP/LTD (Lisman, 1989; Nevian and Sakmann, 2006). The most common assumption on how calcium dynamics are linked to changes of synaptic efficacy is the so-called *calcium-control hypothesis* (Lisman, 1989): high peaks of calcium signal LTP, while prolonged medium-level elevation is the signal for LTD. Plasticity of neocortical synapses between pyramidal cells is traditionally assumed to be presynaptic and in particular to be expressed as a persistent increase/decrease of vesicle release probability (Markram and Tsodyks, 1996). However, more recent studies found also a prominent postsynaptic component (Sjöström et al., 2007), mostly based on alpha-amino3-hydroxy-5-methyl-4-isoxazole propionate receptor (AM-PAR) plasticity, as commonly observed in the hippocampus. Some recent studies (Costa et al., 2017) have investigated the theoretical implications of bi-lateral plasticity and concluded that synapses strive for extreme states (high-conductance high-release probability, low-conductance low-release probability). LTP/LTD are by definition persistent changes, at least in the order of minutes to hours, but presumably longer. This indicates that whatever synaptic state is reached with plasticity, induction needs to be stable at least to some extent. We do not discuss the mechanisms of maintenance per se, as this would require broaching the sub-cellular description level. Instead we are mostly concerned with the number of stable states a synapse can support. Most of the evidence for this topic comes from hippocampal studies. It was shown that postsynaptic (AMPAR-mediated) plasticity is bi-stable and that synapses jump from extreme states of efficacy (Petersen et al., 1998; O’Connor et al., 2005). Presynaptic plasticity instead seems to evolve in a continuous fashion, where synapses can sustain arbitrary levels of efficacy (Enoki et al., 2009).

**Figure 1:**
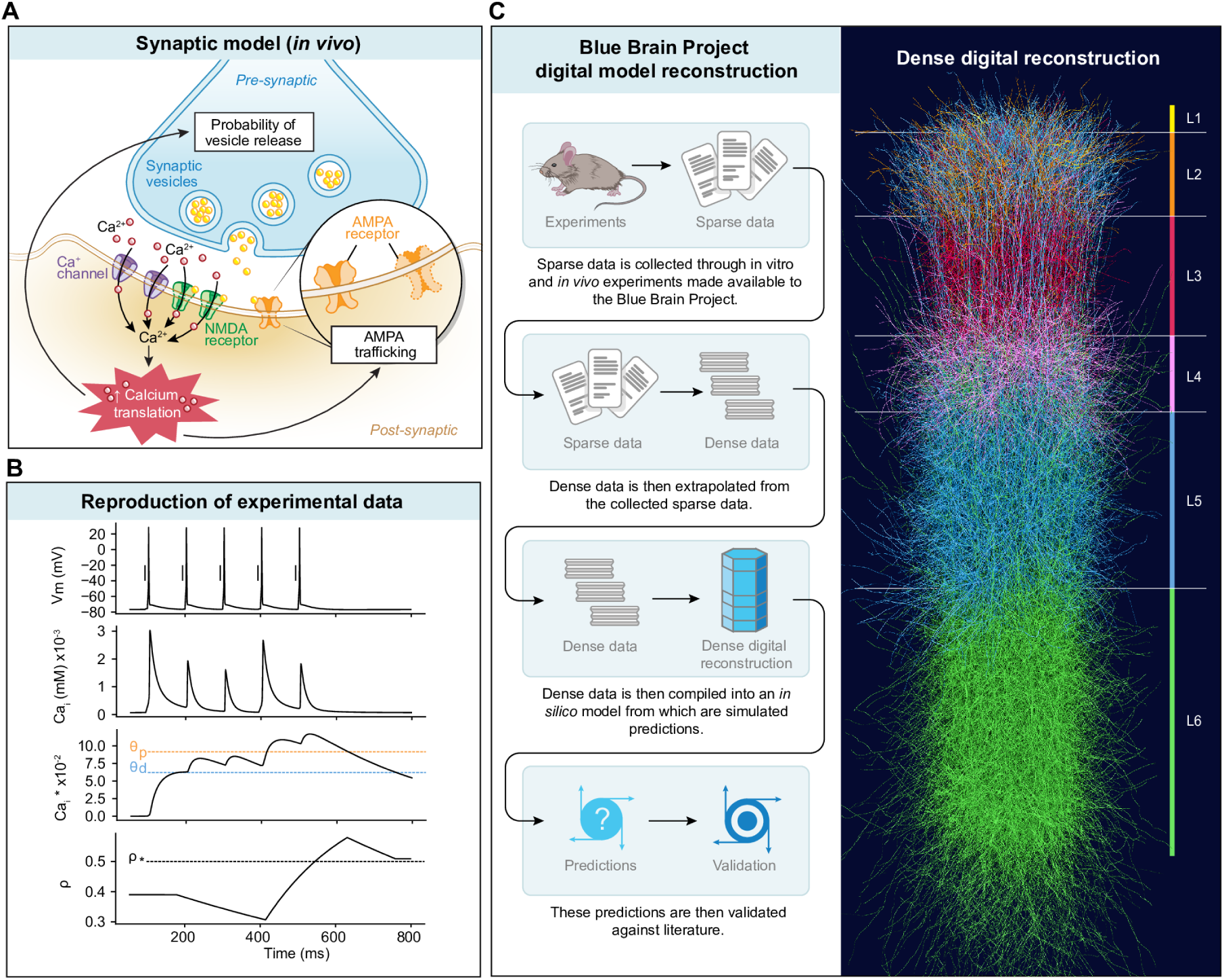
Modeling LTP / LTD in a digital reconstruction of neocortical tissue. **A** Basic model of LTP / LTD dynamics at excitatory synapses. Pre-synaptic vesicle release and subsequent post-synaptic depolarization results in calcium influx in dendritic spines via NMDARs and VDCCs. Calcium signaling activates independent biochemical pathways, leading to long-term changes in AMPAR conductance (postsynaptic expression mechanisms) and/or vesicle release probability (presynaptic expression mechanisms). **B** Evolution of main model variables during coincident activation of pre- and postsynaptic neurons. Presynaptic spikes (vertical dashes) and postsynaptic voltage (solid line) induce postsynaptic calcium transients. The instantaneous calcium concentration is integrated over time to produce a suitable readout for plastic changes. Crossing the depression threshold *θ*_*d*_ causes a drop of the synaptic efficacy *ρ* whereas crossing the potentiation threshold *θ*_*p*_ results in an increase. The net effect of the stimulation protocol is an increase of synaptic efficacy *ρ*. **C** Digital model of neocortical tissue. A standardized process for experimental data integration and generalization is used to generate a dense reconstruction of neocortical circuits. The final model is biologically accurate at multiple levels, from the cellular composition of individual cortical layers to the synaptic innervation patterns of each connection type in the reconstruction.

Based on these findings and assumptions, we defined a simplified model of synaptic plasticity for neocortical synapses. Our model is an extension of previous work by Graupner and Brunel (2012) (see Supplementary Information A.1 for the full description). In brief, the interplay of pre- and postsynaptic activity causes calcium influx in the spine, possibly leading to the activation of LTP/LTD mechanisms if the corresponding calcium thresholds are crossed (Figure 1B). Spine calcium dynamics are modeled as:

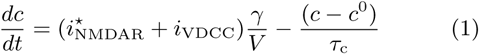

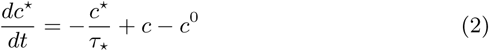

where *c* is the free calcium concentration in the spine head; 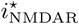 is the calcium component of the NMDAR-mediated current; *i*_VDCC_ is the VDCC-mediated current; *γ* is the fraction of free (non buffered) calcium; *V* is the spine volume; *c*^0^ is the intracellular calcium concentration at rest; *τ*_c_ is the time constant of calcium transients. Equation 2 describes *c*^⋆^, a leaky calcium integrator with time constant *τ*_⋆_. We used *c*^⋆^ as readout for plastic changes in the formalism proposed by Graupner and Brunel (2012):

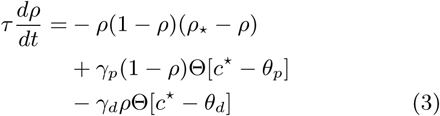

where *ρ* represents the synaptic “efficacy”, with time constant *τ* ; *ρ*_⋆_ delimits the basins of attraction of the two stable states; Θ is the Heaviside function; *θ*_*d*_ and *θ*_*p*_ are respectively the depression and potentiation thresholds; *γ*_*d*_ and *γ*_*p*_ the depression and potentiation rate. The synaptic efficacy *ρ* is dynamically converted into a release probability *u* and AMPAR conductance *g*:

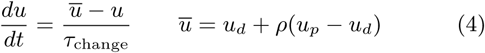

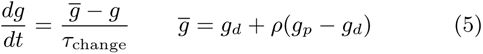

where *u*_*d*_ (*g*_*d*_) and *u*_*p*_ (*g*_*p*_) are the values of the depressed and potentiated state respectively. For simplicity we assumed that these two synaptic variables evolve together, although a more detailed description could be envisioned in the future.

We integrated the modified Graupner and Brunel (2012) model into a digital reconstruction of neocortical circuits (Markram et al., 2015) (Figure 1C). We used this virtual tissue model as the source of neurons and connections for this study. The main advantage is that we could focus on the plasticity aspects only as the synapse and neuron models are already parameterized and constrained to a vast body of experimental evidence. In particular, synaptic innervation is quite accurate in terms of synapse location on the dendrites, number of synapses per connection and vesicle release dynamics (Markram et al., 2015; Reimann et al., 2015; Barros-Zulaica et al., 2019).

To find the parameters for the model we simulated in silico the in vitro experiments performed by Markram et al. (1997b); Sjöström and Häusser (2006), including the known biases of in vitro methods (see Supplementary Information A.2). In brief, pairs of connected L5–L5 (Markram et al., 1997b) and L23–L5 (Sjöström and Häusser, 2006) pyramidal cells (PCs) were stimulated by pairing the activity of the pre- and postsynaptic neurons at various frequencies and time offsets. The mean excitatory post-synaptic potential (EPSP) amplitude was assessed during the minutes preceding the induction protocol and then monitored for 40 minutes after the induction. The last minutes EPSPs were used to assess the amplitude change with respect to baseline. We then used an evolutionary strategy to optimize the model to reproduce the population statistics reported in Markram et al. (1997b); Sjöström and Häusser (2006) (see Supplementary Information A.3). To limit the computational cost of the optimization, we only used 5 out of 9 in vitro experimental protocols, leaving the rest for validation. We assumed that all model parameters but thresholds were constant at all synapses. Potentiation and depression thresholds were instead tailored for every synapse by using a generative model. That is, instead of fitting thresholds for every possible synapse, we assumed that the actual thresholds would be a simple linear combination of the calcium peak during an EPSP *C*_pre_ and the one during a backpropagating action potential (bAP) *C*_post_. These two variables were successfully used in the original Graupner and Brunel (2012) model to fit a generic connection, here we made them synapse specific.

## Results

Our first objective was to evaluate the capability of the model to generalize to novel stimulation protocols. We then tested the model on the two connection types used for parameter estimation, using an unseen set of connections and a larger set of stimulation protocols (L5 PC – L5 PC, Figure 2 A – F; L23 PC – L5 PC, Figure 2 G – L). The relationship between stimulation frequency and timing demonstrates several of the known properties of LTP/LTD at L5 thick-tufted pyramidal cell (TTPC) connections (Figure 2C). The model correctly reproduced LTP frequency dependence as observed in vitro (Markram et al., 1997b; Sjöström et al., 2001), despite the fact that the 20 Hz and above stimulation protocols were not used during training (Fig. 2D). Even more interestingly, we also observed the transition from LTD to LTP at high-frequency stimulation characterized by Sjöström et al. (2001) in V1 slices for the same connection type (Fig. 2E). Spike-timing dependent plasticity (STDP) at 10 Hz stimulation was also matching the available experimental evidence (Fig. 2F), as expected from the good results in training. By extending the STDP time window, we found a characteristic depression-potentiation-depression curve reported in vitro by Sjöström et al. (2001), although at a slightly higher stimulation frequency (20 Hz). The model also captures the marked decrease of LTP levels at L23 PC – L5 PC connections with respect to L5 PC – L5 PC connections (Fig. 2I). In particular, 50 Hz stimulation produces milder LTP at these synapses (Figure 2J). LTD and STDP maintain the same qualitative behavior observed at L5 PC – L5 PC connections, but at lower magnitude (Figure 2 K and L).

**Figure 2:**
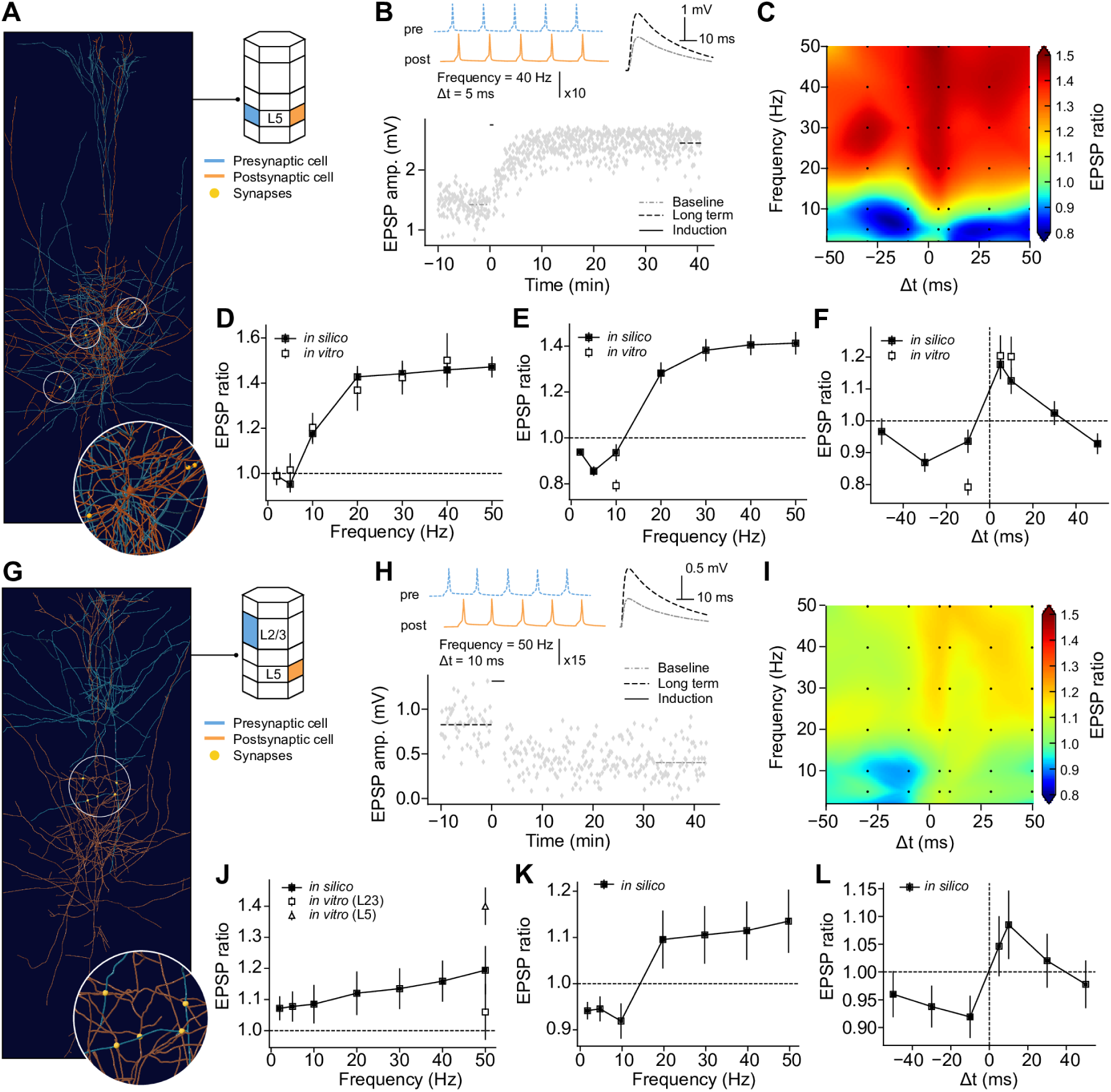
LTP / LTD dynamics in L5-TTPC. **A** Exemplar *in silico* connection between L5-TTPCs. **B** EPSP evolution during plasticity induction experiment (bottom). Induction protocol (top left) consists of 10 bursts of 5 spikes at a frequency of 40 Hz. The presynaptic spike train precedes the postsynaptic one by 10 ms (Δ*t* = +10 ms). Presynaptic spikes depicted for illustration purposes only, see Supplementary Information A.2.1 for details on the neuron simulation methods. Connection strength variation (top right) measured as the ratio of the mean EPSP amplitude after the induction protocol (long term) and the one recorded before the induction protocol (baseline). **C** EPSP ratio heat map for the protocol described in **B**, varying frequency of spike trains and time difference, for a random sample of L5-TTPCs to L5-TTPCs connection (*n* = 100). Experiment configurations marked as black symbols, cubic interpolation elsewhere. **D** Comparison of LTP frequency dependence at Δ*t* = +5 ms *in silico* and *in vitro*. **E** Comparison of LTD frequency dependence at Δ*t* = −10 ms *in silico* and *in vitro*. **F** Comparison of STDP at a frequency of 10 Hz *in silico* and *in vitro*. **G** – **L** as in **A** – **F** for layer 2/3 to layer 5 PC connections. Population data reported as mean ±SEM. For the full distribution of experiment outcomes see SI B.1 (**A** – **F**) and B.2 (**G** – **L**).

Model parameters estimated for L5 TTPC were then applied without modification to other pyramidal connection types from independent in vitro data sources. We observed good agreement between model and experimental data in almost all conditions tested. Typical STDP curves found at L4 PC to L23 PC (Rodríguez-Moreno and Paulsen, 2008) emerge also in our model, including simple pharmacology experiments (Figure 3 A – C). We simulated loading the presynaptic patching pipette with the NMDAR blocker MK801. As we do not explicitly model presynaptic NMDAR, we emulated the effects of MK801 by drastically reducing the LTD rate (see Supplementary Information A.2). Potentiation of L23 PC connections after pairing at 20 Hz stimulation (Egger et al., 1999) is qualitatively captured by our model (Figure 3 D – E). The large uncertainty of the EPSP ratio observed at 20 Hz stimulation is due to a few outlier connections, showing extreme potentiation. To further test the generalization properties of the model, we also considered a set of experiments on layer 4 spiny stellate connections (Egger et al., 1999). Pairing-induced plasticity at this connection type has been shown to be non-NMDAR dependent and, in general, to have quite different mechanisms from pyramidal connections. As expected, our model does not reproduce the *in vitro* results, suggesting that indeed a common set of mechanisms is shared among pyramidal connections only (Figure 3 G – I).

**Figure 3:**
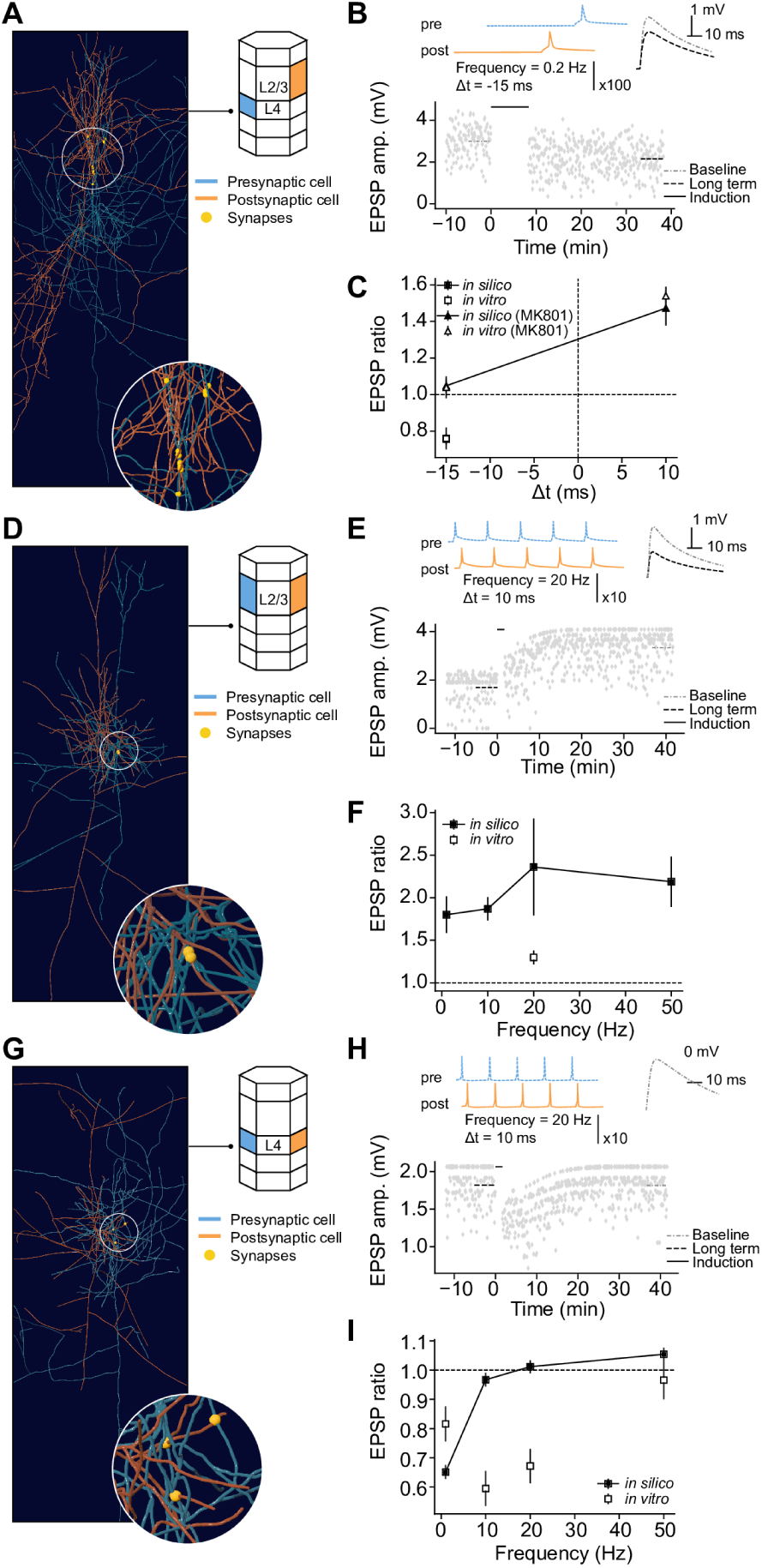
Connection-type specific LTP and LTD. **A** Exemplar *in silico* connection between layer 4 to layer 2/3 PCs. **B** EPSP evolution during plasticity induction experiment (bottom). Induction protocol (top left) consists of 100 pair of spikes every 10 s. The presynaptic spike train follows the postsynaptic one by 10 ms (Δ*t* = −10 ms). Presynaptic spikes depicted for illustration purpose only, see Supplementary Information A.2.1 for details on the neuron simulation methods. Connection strength variation (top right) measured as the ratio of the mean EPSP slope after the induction protocol (long term) and the one recorded before the induction protocol (baseline). **C** Comparison of STDP *in silico* and *in vitro*. **D** – **F** as in **A** – **C** for layer 2/3 to layer 2/3 PC connections. **G** – **I** as in **A** – **C** for layer 4 to layer 4 SSC connections. Population data reported as mean *±* SEM. For the full distribution of experiment outcomes see SI B.3, B.4 (**A** – **C**); B.5 (**D** – **F**) and B.6 (**G** – **I**).

The main advantage of the modeling approach presented here is the possibility to study novel connection types and experimental conditions. To demonstrate that, we provide two examples. First, we predicted the LTP/LTD dynamics of the layer 4 to layer 6 synapses (Figure 4 A – F). As layer 6 pyramidal cell dendrites often terminate in layer 4, it has been proposed that these neurons could associate inputs arriving at the granular and subgranular layers, similarly to what layer 5 cells do with inputs arriving in layer 1 (Ledergerber and Larkum, 2010). Furthermore, the connection between L4 star pyramidal and L6 pyramidal cells is one of the final stages of the corticothalamic feedback loop, with layer 6 cells in turn sending their axons to the thalamus (Qi and Feldmeyer, 2016). For these reasons, it would be particularly interesting to study how synaptic plasticity of layer 4 efferent could modulate the output of layer 6 pyramidal cells. However, the specific LTP/LTD dynamics of this connection type were not yet characterized experimentally, to our knowledge. We found a mild form of LTP frequency dependence very similar to the one observed at the traditional layer 5 TTPC connections (Figure 4D). LTD instead was almost completely absent in our data (Figure 4E). STDP at 10 Hz also shows no LTD, but a graded form of LTP (Figure 4D). Finally, we studied the well known layer 5 TTPC connections at low extracellular calcium concentration, as found *in vivo* (Figure 4 G – L). *In vitro* experiments are traditionally performed at 2 mM calcium, while the *in vivo* levels are estimated to be as low as 1.2 mM. After lowering extracellular calcium (see Supplementary Information A.2), we found that LTP is completely absent (Figure 4I). At high-frequencies, stimulation seems to support only mild LTD (Figure 4 J and K). Time difference sensitivity is also absent in this set of experiments (Figure 4L).

**Figure 4:**
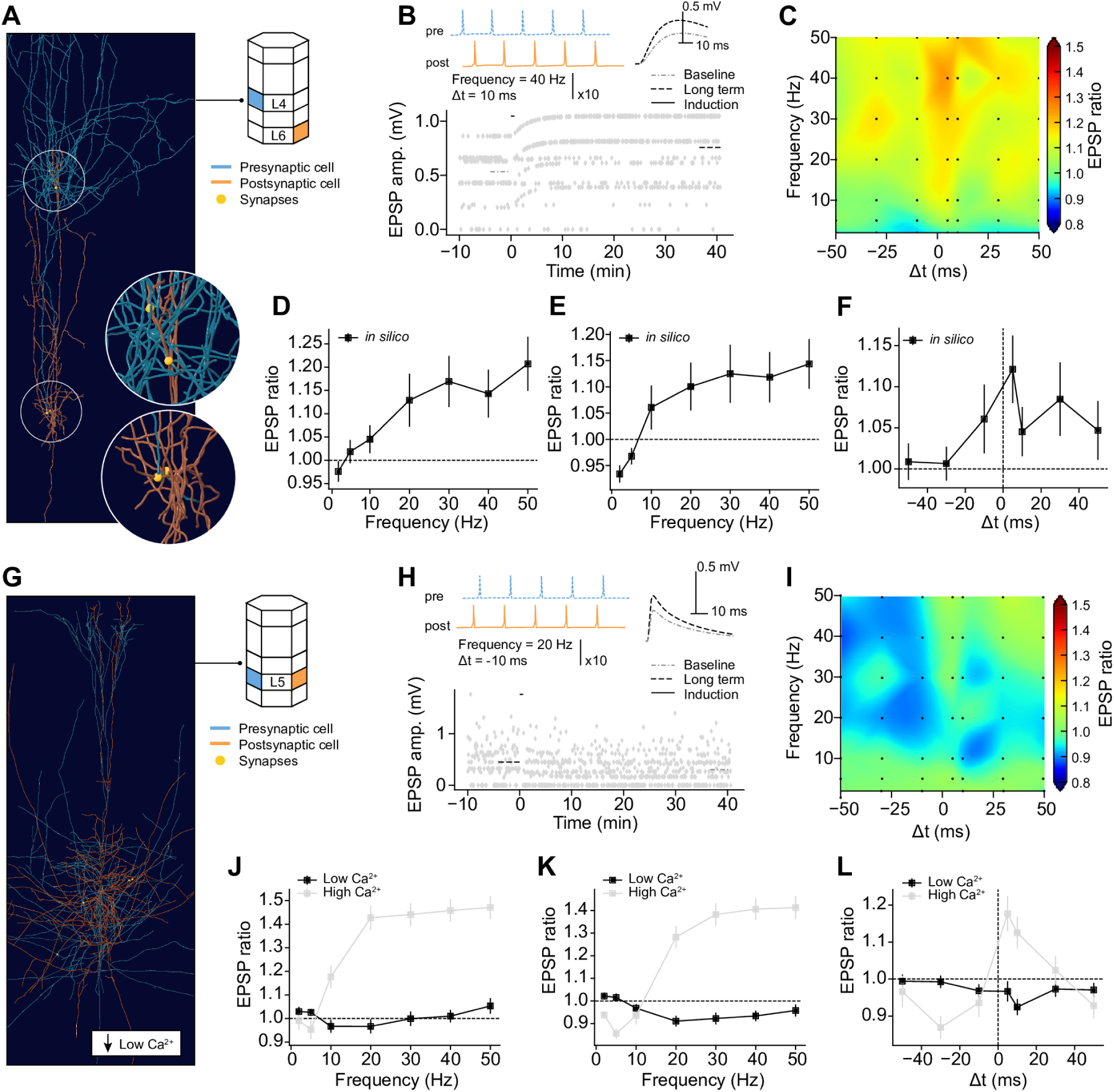
Model predictions. **A** Exemplar *in silico* connection from layer 4 to layer 6 PCs. **B** EPSP evolution during plasticity induction experiment (bottom). Induction protocol (top left) consists of 10 burst of 5 spikes at a frequency of 40 Hz. The presynaptic spike train precedes the postsynaptic one by 10 ms (Δ*t* = +10 ms). Presynaptic spikes depicted for illustration purpose only, see Supplementary Information A.2.1 for details on the neuron simulation methods. Connection strength variation (top right) measured as the ratio of the mean EPSP amplitude after the induction protocol (long term) and the one recorded before the induction protocol (baseline). **C** EPSP ratio heat map for the protocol described in B, varying frequency of spike trains and time difference, for a random sample of layer 5 TTPCs to layer 5 TTPCs connection (*n* = 96–100). Experiment configurations marked as black symbols, cubic interpolation elsewhere. **D** Comparison of LTP frequency dependence at Δ*t* = +5 ms *in silico* and *in vitro*. **E** Comparison of LTD frequency dependence at Δ*t* = −10 ms *in silico* and *in vitro*. **F** Comparison of STDP at a frequency of 10 Hz *in silico* and *in vitro*. **G** – **L** as in **A** – **F** for layer 5 to layer 5 PC connections in low and high extracellular calcium concentration. High calcium data as in Figure 1. Population data reported as mean ± SEM. For the full distribution of experiment outcomes see SI B.7 (**A** – **F**) and B.8 (**G** – **L**).

## Conclusions

In this work we argue that synaptic physiology, cell morphology and innervation patterns govern LTP/LTD dynamics. First we showed that a simple generative model for potentiation and depression thresholds could compensate for the variability of synaptic parameters in the same connection type. That is, despite a variable number of receptors, channels or vesicles, the same model of plastic changes applies to all synapses in a connection. This last result suggests that only a few targeted experiments could be required to characterize the behavior of all other pyramidal connections. Finally, we showed how the model could be used to investigate novel connection types and experimental conditions.

## Discussion

The goal of this study was to show that a single model could capture the plastic behavior of pyramidal cells in the somatosensory cortex. We do not claim nor think that the available experimental data is sufficient to completely understand these forms of plasticity. Instead, we argue that an integrative approach such as the one of this study maximizes the value of the few experimental data points available, it helps homogenizing the results and detecting anomalies. An in-depth analysis of the minimal set of stimulation protocols required to fully characterize all pyramidal cell connections is beyond the scope of this work, as well as meta parameters optimization in the fitting procedure.

This study has several limitations that would need to be addressed in the future. We only consider NMDAR-dependent forms of LTP/LTD at pyramidal cells connections. Although this family includes the vast majority of connections in the neocortex, inhibitory plasticity and other connection types are still likely to play a major role in learning and memory processes. Furthermore, we did not investigate in depth the locus of expression of synaptic changes. We assumed that pre- and postsynaptic changes would match each other over time and ignored the dynamics of this process. This choice was motivated in part by the lack of experimental constraints and in part by the results of recent theoretical studies (Costa et al., 2017). We obtained only moderated levels of LTD compared to in vitro experiments. Low-pass filtering the calcium concentration inevitably trades part of the STDP accuracy for a better handling of the LTP frequency dependence. A possible solution for this problem would be to decouple the LTP and LTD pathways, as experimented in other recent models. Although this is certainly possible, it would substantially increase the number of free parameters to fit and so the cost of the optimization.

As previously mentioned, many more models of synaptic plasticity exist (for a review see Manninen et al. (2010)). In this work we adapted a previously published calcium-based model (Graupner and Brunel, 2012) to match the description level of our neural tissue model (Markram et al., 2015). This approach allowed us to take into account important aspect of synaptic physiology often neglected in plasticity studies, such as stochastic vesicle release, while limiting the number of free parameters to constrain. Furthermore, a direct dependence on calcium dynamics allowed us to extrapolate the behavior of pyramidal connections at physiological calcium concentration, considerably lower than the canonical 2 mM of in vitro experiments. Our choice of a base model was then motivated by practical considerations. Moreover, several ingredients of the Graupner and Brunel (2012) model are based on well established ideas in the field. For example, the model takes advantage of the full calcium time course, rather than just the peaks, to identify plasticity-inducing events. This general principle has already been exploited more or less explicitly in many other models (Rubin et al., 2005; Clopath and Gerstner, 2010; Ebner et al., 2019). Here we introduced a calcium integrator to associate synaptic events beyond the duration of individual calcium transients and to compensate for synaptic unreliability. The original Graupner and Brunel (2012) model lacks this term as the same purpose could be served by a longer calcium time constant and reliable vesicle release. Regardless of the actual mathematical formalism, simpler or more detailed models could benefit from the parameterization here proposed to match the behavior of different connection types in network studies. Finally we argue that the present study shows the importance of modeling dendrites to study synaptic plasticity as the cellular level. We used morphological neuron models to generalize the optimized parameters to all connection types, leveraging dendritic dynamics to normalize calcium. However, the main reason to model dendrites in plasticity studies is the need to move from the stimulation protocols of *in vitro* experiments to more realistic conditions. In particular, we showed how dropping calcium to *in vivo* levels profoundly limits LTP/LTD dynamics, making traditional plasticity induction protocols largely ineffective. It is nowadays more and more accepted that N-methyl-D-aspartate (NMDA) spikes and other local dendritic events are the real driving signal for plastic changes. STDP protocols and related are a very effective tool to investigate the landscape of synaptic changes and could provide a window into more complex activation dynamics, such as NMDA or calcium spikes, even without triggering them. A model such as the one presented here could allow to study these phenomena with higher specificity and in a more realistic network environment.

## Acknowledgements

The authors thank Michael Hines for helping with synapse model implementation in NEURON; Mariana Vargas-Caballero for sharing NMDAR data; Veronica Egger for sharing in vitro data and for clarifications on the analysis methods; Jesper Sjostrom for sharing in vitro data and discussions; Ralf Schneggenburger for discussion and clarifications on the NMDAR’s calcium current; Francesco Casalegno for discussion on model fitting; Michael Reimann and Max Nolte for feedback on the manuscript; Wulfram Gerstner and Guillaume Bellec for discussions.

## A. Supplementary methods

Part of the text and figures in the following sections are reproduced or adapted from Chindemi (2018).

### A.1. Synapse model

The long-term potentiation (LTP)/long-term depression (LTD) model used in this work was built on top of the basic alpha-amino-3-hydroxy-5-methyl-4-isoxazole propionate receptor (AMPAR) / N-methyl-D-aspartate receptor (NMDAR) synapse described in Markram et al. (2015). For the sake of completeness, we present in the following sections all the components of the synapse model, including those previously developed (Markram et al., 2015). Unless differently stated, model parameters for each connection type are available online at: https://bbp.epfl.ch/nmc-portal/microcircuit.

#### A.1.1. AMPAR

The AMPARs are described using a double exponential conductance profile (Ramaswamy et al., 2012; Markram et al., 2015; Ramaswamy et al., 2015):

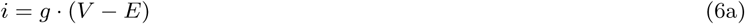

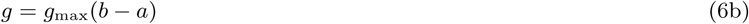

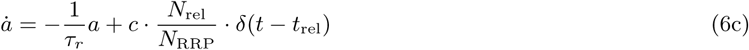

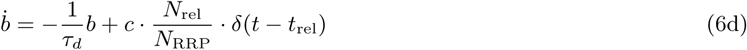

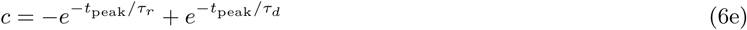

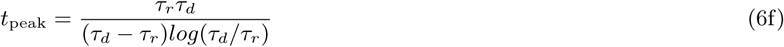

where *i* is the current produced by the synaptic population of AMPAR; *E* is the reversal potential of the receptor; *V* is the membrane potential; *g* is the conductance of the receptor population, with peak *g*_max_; *a* models the rising component of the conductance, with time constant *τ*_*r*_; *b* models the decaying component of the conductance, with time constant *τ*_*d*_; *t*_rel_ is the time of a successful release event; *N*_rel_ is the number of vesicles released at time *t*_rel_ out of *N*_RRP_ available in the readily-releasable pool (RRP) (see A.1.3); *c* is a normalization factor such that when *t* = *t*_rel_ + *t*_peak_, *g* = *g*_max_; *t*_peak_ is the time to peak of the conductance.

The peak AMPAR conductance *g*_max_ is dynamically linked to the synaptic efficacy *ρ* of the long-term plasticity model (see section A.1.7). As the synaptic efficacy is assumed to be bistable (Graupner and Brunel, 2012), so it is the AMPAR conductance:

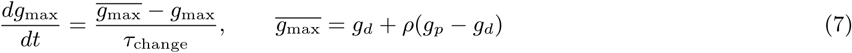

where 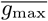 is the target value of *g*_max_; *g*_*d*_ and *g*_*p*_ are the conductance values of the depressed and potentiated state respectively; *τ*_change_ is the convergence time constant. The decision to make *g*_max_ a dynamic variable, as opposed to an instantaneous mapping as in Graupner and Brunel (2012), is dictated by the observation that the expression of synaptic plasticity is known to be slow compared to its induction. It is indeed common to observe a buildup of the plastic changes over the course of several minutes after the induction protocol in vitro (Markram et al., 1997b; Sjöström et al., 2001).

All the parameters of the AMPAR model are prescribed by the tissue model (Markram et al., 2015), with the exception of the novel *g*_*d*_, *g*_*p*_ and *τ*_change_. We decided to fix *τ*_change_ = 100s, based on the typical time course observed in vitro (Markram et al., 1997b; Sjöström et al., 2001). This value could obviously be refined to exactly match experimental dynamics, as done for other parameters in this model, but we did not expect major differences and so avoided to add another parameter to the optimization procedure (see section A.3). It has been shown in the hippocampus that *g*_*p*_ ≈ 2*g*_*d*_ (O’Connor et al., 2005). We therefore initialized each individual synapse as follows:

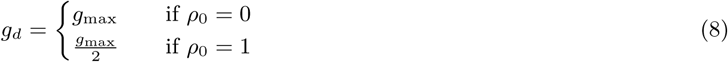

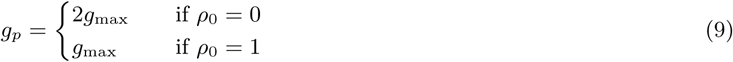

where *ρ*_0_ is the initial value of the synaptic efficacy, *ρ*_0_ = 0 indicates a depressed synapse while *ρ*_0_ = 1 a potentiated one (see section A.1.7 for further details).

#### A.1.2. NMDAR

The NMDARs are described using a double exponential conductance profile with magnesium block dynamics (Ramaswamy et al., 2012; Markram et al., 2015; Ramaswamy et al., 2015).

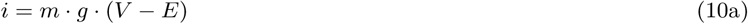

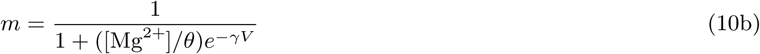

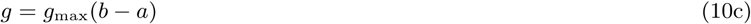

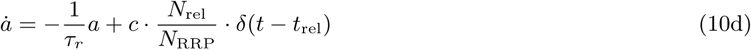

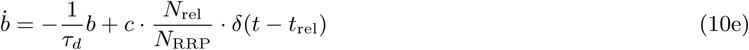

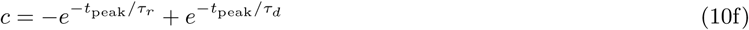

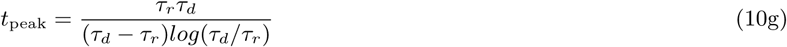

where *i* is the current produced by the synaptic population of NMDAR; *E* is the reversal potential of the receptor; *V* is the membrane potential; *m* is the magnesium block gating variable Jahr and Stevens (1990); *θ* is an appropriate scaling factor of the extracellular magnesium concentration [Mg^2+^]; *γ* is the slope of magnesium voltage dependence; *g* is the conductance of the receptor population, with peak *g*_max_; *a* models the rising component of the conductance, with time constant *τ*_*r*_; *b* models the decaying component of the conductance, with time constant *τ*_*d*_; *t*_rel_ is the time of a successful release event; *N*_rel_ is the number of vesicles released at time *t*_rel_ out of *N*_RRP_ available in the RRP (see A.1.3); *c* is a normalization factor such that when *t* = *t*_rel_ + *t*_peak_, *g* = *g*_max_; *t*_peak_ is the time to peak of the conductance.

The fractional calcium current *P*_*f*_ through the NMDAR can be calculated from the Goldman-Hodgkin-Katz (GHK) flux equation (Schneggenburger et al., 1993):

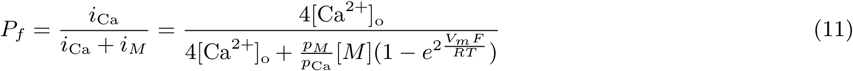

where *i*_Ca_ and *i*_*M*_ are respectively the current due to calcium ions and the one due to all monovalent ions; [Ca^2+^]_o_ is the extracellular calcium concentration; [*M*] is the concentration of monovalent ions, assumed to be identical inside and outside the cell membrane; *p*_Ca_ and *p*_*M*_ are respectively the permeability to calcium ions and the one to monovalent ions; *V*_*m*_ is the membrane voltage; *T* is the temperature; *F* is the Faraday’s constant and *R* is the ideal gas constant.

Unfortunately Equation 11 cannot be directly used in our model as a scaling factor for the total NMDAR current because it loses physical meaning in the limit of *V*_*m*_ → 0 mV, a threshold often crossed during action potential (AP). That is, *P*_*f*_ → 1 for *V*_*m*_ → 0, but as also *i*_NMDAR_ → 0, the total calcium current would be incorrectly estimated. Based on the observation that the calcium current approaches 0 for *V* ≥ 40 mV (Schneggenburger et al., 1993), we approximated the calcium component of the NMDAR conductance *g*_Ca_ as a separate ion channel with reversal potential *E* = 40 mV:

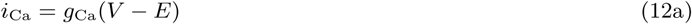

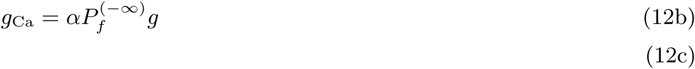

where *α* is an appropriate scaling factor, determined by comparing model simulation results to Equation 11; 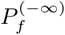 is the fractional calcium current at very negative potentials, all other parameters as in Schneggenburger et al. (1993)); *E* was assumed to be independent of extracellular calcium concentration, at least for values around 1 mM (Jahr and Stevens, 1993). The proposed model was then tested against the findings of Schneggenburger et al. (1993) and accepted as a good approximation in the physiological voltage range and at relevant extracellular calcium contention.

#### A.1.3. Neurotransmission

Vesicle release is described using a stochastic version of the canonical Tsodyks-Markram (TM) model with multi-vesicular release (MVR) (Tsodyks and Markram, 1997; Ramaswamy et al., 2012; Markram et al., 2015; Ramaswamy et al., 2015; Barros-Zulaica et al., 2019). It is effectively analogous to the traditional binomial model of vesicle release *B*(*N*_RRP_, *U* (*t*)), where *N*_RRP_ is the number of vesicles available at any given moment in the RRP and is and *U* (*t*) is the dynamical release probability of the TM formalism:

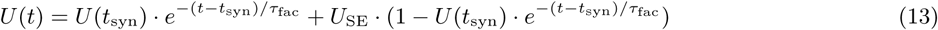

where *U*_SE_ is the limit value of the release probability (i.e. the release probability of a synapse not activated in a very long time); *t*_syn_ is the time of the last presynaptic spike; *τ*_fac_ is the facilitation time constant. Vesicle recovery also follows the binomial model *B*(*N*_R_, 1 − *P*_surv_(*t*)), where *N*_R_ is the number of vesicles awaiting recovery from release and *P*_surv_(*t*) is the survival probability of the un-recovered state:

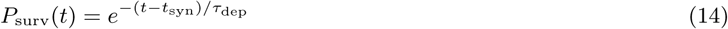

where *τ*_dep_ is the depression time constant.

The limit value of the release probability *U*_SE_ is dynamically linked to the synaptic efficacy *ρ* of the long-term plasticity model (see section A.1.7), as done for the peak AMPAR conductance (see section A.1.1). As we decided to match exactly pre- and post-synaptic changes by having a single efficacy variable *ρ*, also *U*_SE_ was assumed to be bistable:

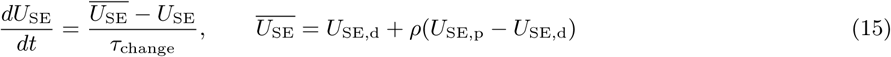

where 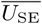 is the target value of *U*_SE_; *U*_SE,d_ and *U*_SE,p_ are the release probability of the depressed and potentiated state respectively; *τ*_change_ is the convergence time constant.

All the parameters of the vesicle release model are prescribed by the tissue model (Markram et al., 2015), with the exception of the novel *U*_SE,d_, *U*_SE,p_ and *τ*_change_. We decided to fix *τ*_change_ = 100s, based on the typical time course observed in vitro (Markram et al., 1997b; Sjöström et al., 2001). This value could obviously be refined to exactly match experimental dynamics, as done for other parameters in this model, but we did not expect major differences and so avoided to add another parameter to the optimization procedure (see section A.3). It has been shown in the hippocampus that presynaptic plasticity is graded and bidirectional (Enoki et al., 2009). For this reason, a model with three stable states or even a continuum of stable states would have been more appropriate to capture long-term changes of release probability. However, it would also have required to decouple pre- and post-synaptic changes, greatly increasing the number of free parameters to constraint. We therefore decided to maintain the bistable hypothesis for this first release of the model and assume saturating potentiation and depression events. That is, we assumed an exponential relationship between the potentiated and depressed state to account for release probability saturation:

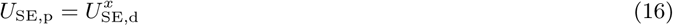

where x is an appropriate constant. From the work of Enoki et al. (2009) we can estimate the value of *x* to match the set of experiments where strong and repeated LTP/LTD was induced. We found that *x* ∈ (0.1, 0.25) would provide a satisfactory approximation and chose *x* = 0.2 for simplicity. We then initialized *U*_SE,d_ and *U*_SE,p_ at every synapse as:

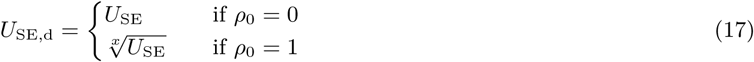

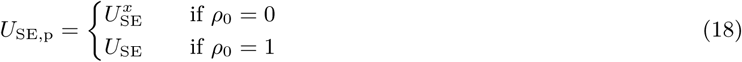

where *ρ*_0_ is the initial value of the synaptic efficacy, *ρ*_0_ = 0 indicates a depressed synapse while *ρ*_0_ = 1 a potentiated one (see section A.1.7 for further details).

#### A.1.4. VDCC

In this work, we only considered high voltage activated (HVA) calcium channels. A simple but inactivating population of R-type voltage-dependent calcium channel (VDCC) was designed in the Hodgkin-Huxley (HH) formalism (Magee and Johnston, 1995; Sabatini and Svoboda, 2000).

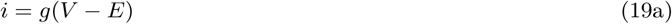

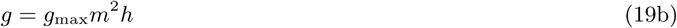

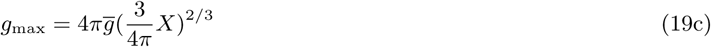

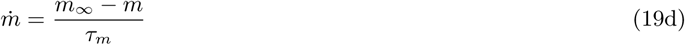

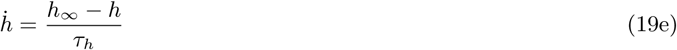

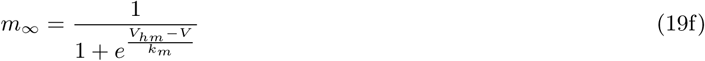

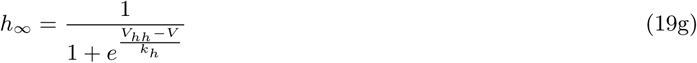

where *i* is the current produced by the channel population; *V* is the membrane potential; *E* is the reversal potential for calcium; *g* is the conductance of the population, with peak *g*_max_, calculated assuming a spherical spine head; 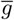 is the VDCC surface area density; *X* is the spine head volume (see A.1.5); *m* is the activation variable, with time constant *τ*_*m*_ and steady state *m*_∞_; *h* is the inactivation variable, with time constant *τ*_*h*_ and steady state *h*_∞_; *V*_*hm*_ is the half-maximum activation voltage and *k*_*m*_ is the slope factor; *V*_*hh*_ is the half-maximum inactivation voltage and *k*_*h*_ is the slope factor. All model parameters were obtained from literature, as summarized in Table 1.

**Table 1:**
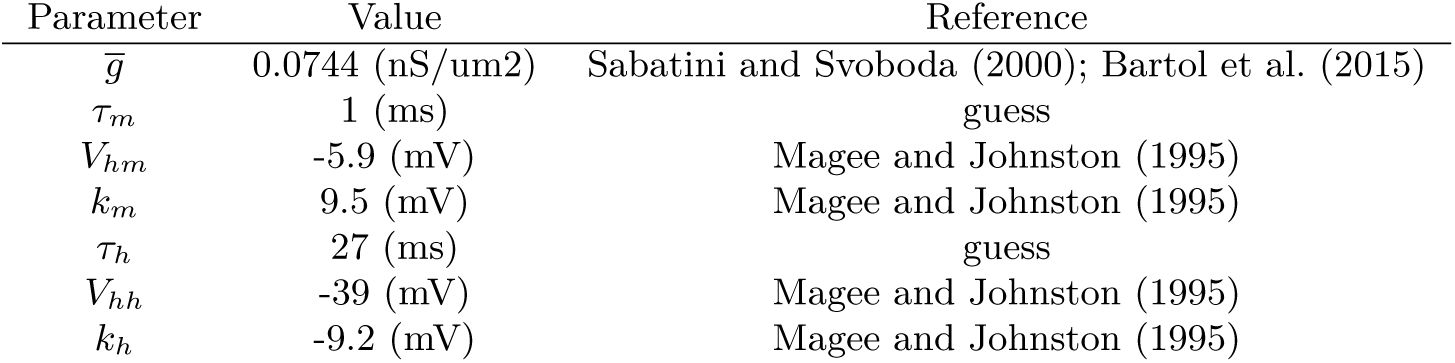
VDCC model parameters.

| Parameter | Value | Reference |
| --- | --- | --- |
| $\bar{g}$ | 0.0744 (nS/um2) | Sabatini and Svoboda (2000); Bartol et al. (2015) |
| $\tau_m$ | 1 (ms) | guess |
| $V_{hm}$ | -5.9 (mV) | Magee and Johnston (1995) |
| $k_m$ | 9.5 (mV) | Magee and Johnston (1995) |
| $\tau_h$ | 27 (ms) | guess |
| $V_{hh}$ | -39 (mV) | Magee and Johnston (1995) |
| $k_h$ | -9.2 (mV) | Magee and Johnston (1995) |

#### A.1.5. Spine

The vast majority of excitatory synapses are located on dendritic spines. With respect to synaptic plasticity, spines are particularly important because they act as biochemical compartments, granting specificity of synaptic changes. Furthermore, spine volume is a key determinant of intracellular calcium concentration. In this work we do not explicitly model spines for computational reasons, but we do account for biochemical compartmentalization and volume-related effects on calcium concentration. Specifically, a spherical spine head is assumed for all computations and calcium ions are not allowed to diffuse into the parent dendrite. Spine volume parameterization was carried out using data from electron microscope (EM) reconstructions. A crucial aspect of the parameterization was the correlation introduced between spine volume and NMDAR conductance (Arellano et al., 2007). This feature allowed to generate VDCC conductance proportional to NMDAR for every synapse (see Section A.1.8).

#### A.1.6. Postsynaptic calcium dynamics

The postsynaptic calcium model was designed combining data from several experimental and theoretical sources (Sabatini et al., 2002; Cornelisse et al., 2007). In brief, calcium current can flow through cell membrane via two paths: NMDARs and VDCCs. After entering the spine, calcium ions quickly bind to endogenous buffers and only a small fraction remains free. Slower mechanisms, such as calcium pumps and diffusion, are in place to re-establish the intracellular calcium concentration. These dynamics are all summarized in a single ordinary differential equation (ODE), modeling free calcium at the postsynaptic membrane surface:

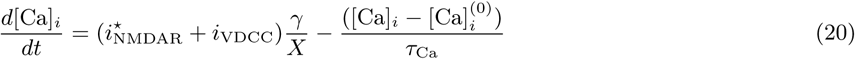

where 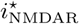 is the calcium component of the NMDAR-mediated current, as later described; *i*_VDCC_ is the VDCC-mediated current; *γ* is the fraction of free (non buffered) calcium; *X* is the spine volume; 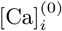 is the intracellular calcium concentration at rest; *τ*_Ca_ is the time constant of calcium transients.

Intracellular calcium dynamics are key for the induction of long-term synaptic changes and so they have been heavily studied in the context of spike-timing dependent plasticity (STDP) (Nevian and Sakmann, 2006). Even tough it is nowadays clear that calcium is the main driver for such changes, how exactly synapses could *read* this signal is still debated. One of the most established theories is the so called *calcium control hypothesis*: LTP is induced by large calcium transients, while LTD is triggered by prolonged medium-level transients. Although this simple explanation is very elegant and appealing from the theoretical point of view, its experimental validation has proven to be particularity tricky and many open questions remain. In particular, it was shown that meditum- to high-frequency stimulation can trigger LTP while low-frequency does not (Markram et al., 1997b), but whether the former causes calcium accumulation at the synapse is unknown (Manninen et al., 2010). Some experimental evidence seems to suggest that small-capacity and fast-binding calcium reaction partners need to be saturated before the mechanisms responsible of LTP and LTD could be activated (for a review see (Manninen et al., 2010)). Indeed preliminary analysis showed that calcium does not substantially accumulate in our model at medium- to high-frequency stimulation. For this reason, we speculate that the frequency dependence of LTP could not be explained based on free-calcium only and requires to consider the different calcium reaction partners. To account for such dynamics without explicitly modeling the endogenous buffers, we introduced the calcium integrator 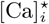:

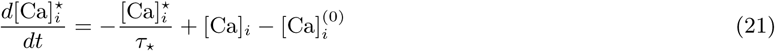

where *τ*_⋆_ is an appropriate time constant for calcium integration, fitted to match experimental data on LTP/LTD (see Section A.3). Boundaries for this value were chosen considering both the average duration of an NMDAR-mediated spike (Schiller et al., 2000), considered the key signal for triggering LTP, and the typical stimulation window of LTP-inducing protocols (Markram et al., 1997b). The idea behind the introduction of 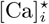 is to prevent a single and isolated calcium transient from inducing any long-term change of synaptic efficacy, while allowing several pulses at high frequency, such as those used in STDP protocols, or large cooperative events, such as NMDAR-mediated spikes, to produce enough activation to trigger a long-term synaptic change. The calcium control hypothesis, in the form of the model proposed by Graupner and Brunel (2012), was than applied to the 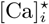 signal rather than the free calcium [Ca]_*i*_ (see Section A.1.7).

#### A.1.7. Long-term plasticity

In this work, we adapted the calcium-based model of STDP by Graupner and Brunel (2012) to account for more realistic synaptic dynamics. The mathematical formalism was left mostly unchanged. As in the original work by Graupner and Brunel (2012), a support variable *ρ* describes the current state and variations of synaptic efficacy:

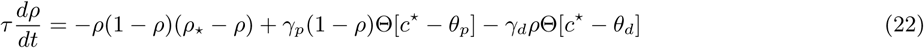

where *τ* is an appropriate time constant; *ρ*_⋆_ delimits the basin of attraction of the two stable states; Θ is the Heaviside function; *θ*_*d*_ and *θ*_*p*_ are respectively the depression and potentiation thresholds; *γ*_*d*_ and *γ*_*p*_ the depression and potentiation rate; *c*^⋆^ is the calcium integrator described in Section A.1.6. The synaptic efficacy *ρ* is dynamically converted into an AMPAR conductance *g* and a release probability *u*, as described respectively in Section A.1.1 and Section A.1.3. The synaptic noise term, present in the original Graupner and Brunel (2012) model, was removed here as we already account for several stochastic aspect of synaptic transmission and membrane potential fluctuations in the synapse and neuron models.

#### A.1.8. Synaptic parameters correlation

It is well established that several morphological variables of excitatory synapses are correlated. Several studies have shown that postsynaptic density (PSD) area is strongly correlated with pre- and post-synaptic variables, such as spine head volume (Harris and Stevens, 1989; Schikorski and Stevens, 1999; Arellano et al., 2007), bouton volume and number of vesicles (Harris and Stevens, 1989). Such findings suggest that a certain degree of parameter correlation is required/beneficial for a synapse model. In particular, wherever correlations are particularly strong, the value of unknown parameters could be well predicted from the available ones. Furthermore, correlations might help us and nature to normalize synaptic dynamics (i.e. calcium transients) and so simplify the task of applying the same recipe (i.e. calcium thresholds for LTP/LTD) to a large set of heterogeneous synapses. Based on the available experimental evidence, we imposed the following correlations to synaptic model variables. We are assuming mathing pre- and post-synaptic changes, so a high correlation between release probability and conductance is implicit. Unfortunately we are not aware of any report explicitly quantifying this quantity, so we chose a generic high value to be refined in the future:

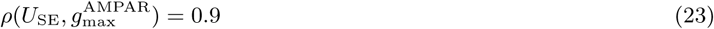

The relationship between total number of vesicles and PSD area / spine volume was estimated by Harris and Stevens (1989). Assuming the total number of vesicles is a good proxy for the size of the RRP and that the PSD area is so for conductance, we imposed:

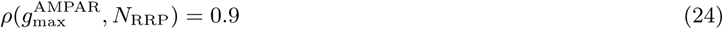

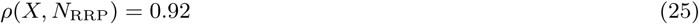

Finally, from Arellano et al. (2007) we sat the correlation between spine volume and synaptic conductance. This correlation was particularly precious because it allowed us to predict spine volumes, an unknown variable, from conductance, a constrained parameter in the tissue model (Markram et al., 2015) (see Section A.1.5):

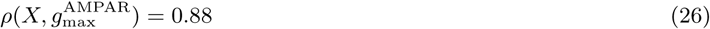

A valid correlation matrix needs to be positive-semidefinite. We filled the missing entries using a simple algorithm proposed by Kahl and Günther (2008), obtaining the final correlation matrix *M* used in this work for the vector of parameters *P* :

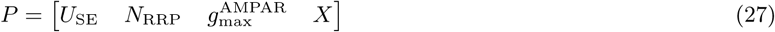

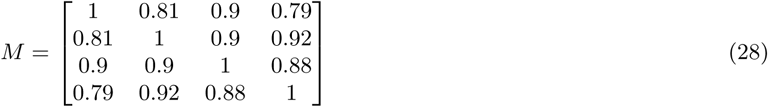

### A.2. In silico electrophysiology

#### A.2.1. Simulation

All in silico experiments were performed using NEURON (Hines and Carnevale, 1997) and the Blue Brain Project (BBP) tissue model (Markram et al., 2015). The only modification applied to the electrical model of digital cells was the removal of calcium-activated potassium conductance from the cell body and axon initial segment. This change was required to eliminate extreme hyper-polarization during high frequency stimulation, an artefact expected to be corrected in future releases of the tissue model (Markram et al., 2015). As in NEURON synapses are considered postsynaptic processes activated by a presynaptic trigger, we did not need to simulate the presynaptic cell during our experiments and we would not obtain any benefit in doing so. Rather we computed the desired spike timing and fed it to the synaptic processes, hosted on the postsynaptic cell. To further reduce the computational cost of each simulation, we fast-forwarded the convergence of the LTP/LTD model variables after the induction. That is, after establishing whether stimulation was sufficient to cause a state change, we moved the interested synaptic variables to their new fixed points rather than simulating their (slow) convergence. This trick was used during model fitting to substantially reduce the computational cost, while disabled for experiments showing the full convergence dynamics. Fast-forwarding the convergence dynamics does not affect our results in any way. Calcium thresholds cannot be reached during the convergence phase by definition, as they are always higher than the transients generated by sparse pre- and post-synaptic stimulation (see Section A.3). The only difference between a regular simulation and a fast-forwarded one would be due to the random number generator (RNG), as in the former case excitatory post-synaptic potentials (EPSPs) are also generated during the convergence phase.

#### A.2.2. Reproducing in vitro experiments

**Figure A.1:**
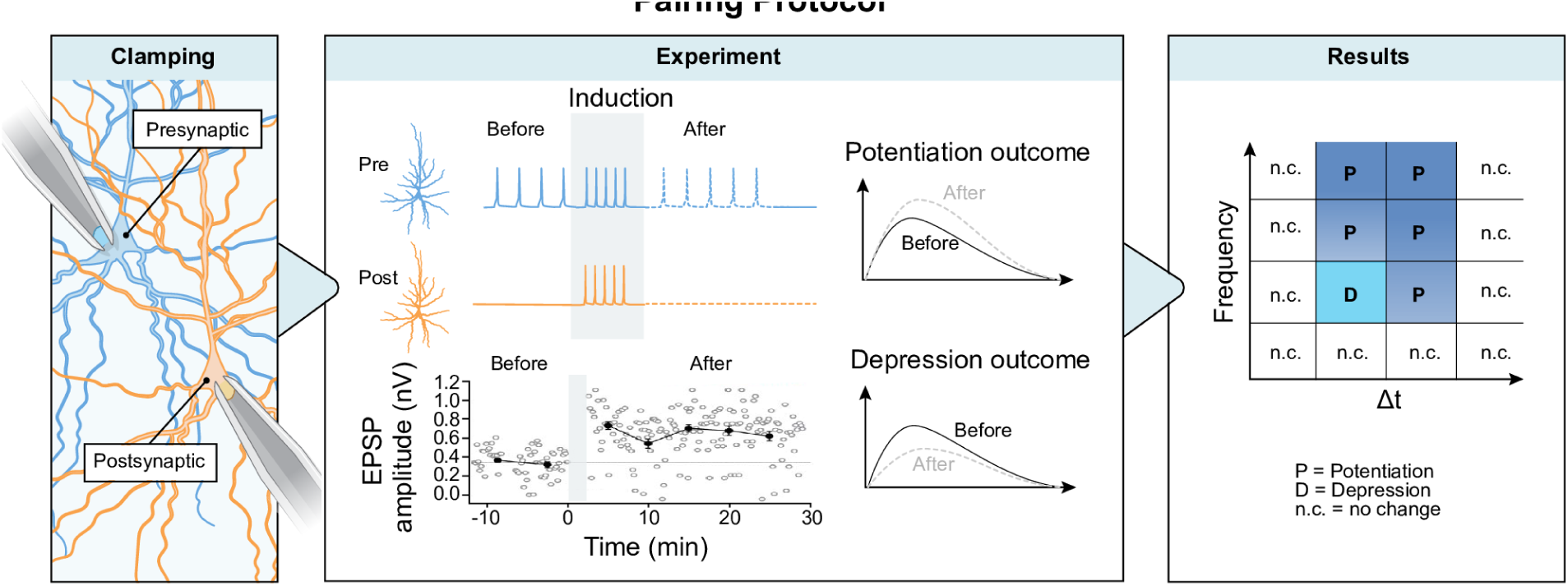
A typically paired in vitro experiment. The protocol includes three phases: (a) an initial assessment of the connection strength; (b) a plasticity-induction protocol where the activity of the two neurons is paired at a given frequency and time difference; (c) a prolonged evaluation of connection strength.

We attempted to digitally reproduce multi patch-clamp in vitro experiments as close as possible, including their typical biases. Neurons were randomly selected mimicking the tendency of experimenters to patch nearby cells on the same focal plane (i.e. 50 × 50 x 10 µm volume for layer 5 thick-tufted pyramidal cell (TTPC) connections). This typical bias is motivated by the desire of maximizing connection probability in lab experiments and so slice yield. We only considered paired in vitro recordings, as they provide the most reliable and controlled source of experimental evidence on LTP/LTD (see Figure A.1). The specifics of individual experiments for each connection type considered in this work are described in the following paragraphs.

##### Layer 5 PC to Layer 5 PC connections

We considered the methods in Markram et al. (1997b); Sjöström and Häusser (2006). Connections were selected from random volumes of 50 × 50 x 10 µm. Both stimulation protocols and data analysis could be reproduced almost exactly.

##### Layer 23 PC to Layer 5 PC connections

We considered the methods in Sjöström and Häusser (2006). Connections were selected from random volumes of 50 × 700 x 10 µm. Both stimulation protocols and data analysis could be reproduced almost exactly.

##### Layer 4 PC to Layer 23 PC connections

We considered the methods in Rodríguez-Moreno and Paulsen (2008). Connections were selected from random volumes of 50 x *max* x 10 µm. Both stimulation protocols and data analysis could be reproduced almost exactly. We emulated the effects of MK801 by reducing tenfold the LTD rate:

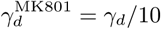

##### Layer 23 PC to Layer 23 PC connections

We considered the methods in Egger et al. (1999). Connections were selected from random volumes of 50 × 50 x 10 µm. The specific somatosensory cortex region of the in vitro experiments was different from the one of our tissue mode (barrel cortex in Egger et al. (1999), non-barrel cortex in (Markram et al., 2015)). Data analysis was performed following the directions in Egger et al. (1999). However, the Gaussian weighting method could not be applied to our data. This technique, used to reduce the error on the mean EPSP estimate, has several requirements on data distribution and correlations. In particular, the mean EPSP amplitudes must have a Gaussian distribution. This condition is not always met by our data after plasticity induction. For this reason, great care has to be taken to correctly compare the results of our model with the *in vitro* data in Egger et al. (1999). This result is not surprising, considering that we did not impose any constraints on the distribution of EPSP amplitudes after plasticity during model fitting, nor invalidating, as we just seek qualitative agreement of the protocol results (i.e. LTP vs LTD or no change).

##### Layer 4 SSC to Layer 4 SSC connections

We considered the methods in Egger et al. (1999). Connections were selected from random volumes of 50 × 50 x 10 µm. Same considerations for layer 23 PC to layer 23 PC connections apply here.

##### Layer 4 PC to Layer 6 PC connections

We adapted the stimulation protocols and analysis from Markram et al. (1997b). Connections were selected from random volumes of 50 × 800 x 10 µm. Postsynaptic neurons spontaneously spiking during EPSP amplitude assessment were excluded from analysis.

##### Layer 5 PC to Layer 5 PC connections in low calcium

We considered the methods in Markram et al. (1997b); Sjöström and Häusser (2006). Connections were selected from random volumes of 50 × 50 x 10 µm. We model the low calcium conditions in vivo by (a) reducing the synaptic release probability to 15% of its in vitro value (Markram et al., 2015); (b) adapting the calcium reversal potential; and (c) recomputing the fractional component of calcium current through the NMDARs.

### A.3. Model fitting

Before describing the optimization procedure, we need to clarify our strategy to obtain calcium thresholds for a pool of heterogeneous synapses. That is, in our tissue model every synapse is unique as its parameters are randomly drawn from appropriate distributions (Markram et al., 2015). Furthermore, synapses are spatially distributed on complex dendritic trees. For these reason, each synapse shows diffident calcium dynamics and no fixed thresholds could apply to all of them, even withing the same connection type. To solve this problem, we assumed a generative model where plasticity thresholds are expressed as linear combinations of the calcium readout peaks during isolated presynaptic activation *C*_pre_ and isolated postsynaptic activation *C*_post_:

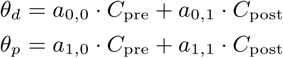

where *a*_*i,j*_ are appropriate constants obtained during model fitting. The idea behind this solution is that synapses are likely to self-regulate calcium threshold using appropriate homeostatic mechanisms. *C*_pre_ and *C*_post_ were already part of the original Graupner and Brunel (2012) formalism as free parameters. Here instead they are measured quantities used to customize thresholds for each synapse. In our preliminary attempts to fit the model, we found that using a separate set of *a*_*i,j*_ parameters for apical and basal synapses provided better results. We do not exclude that using a nonlinear combination of *C*_pre_ and *C*_post_ could allow to unify apical and basal thresholds, but we have not explored this possibility yet. All other parameters were identical for apical and basal synapses.

To fit the long-term plasticity model we then needed to find appropriate values for a limited set of free parameters (see Table 3). To further reduce the size of the search space, we decided to exclude from this list any parameter that could be fixed from Graupner and Brunel (2012) or from test runs. For example, the two time constants *τ* and *τ*_change_ were estimated from preliminary fitting attempts and are compatible with previously used parameter sets (see Supplementary Information in Graupner and Brunel (2012)). Values for fixed parameters are reported accordingly in Section A.1. To fit the remaining parameters we considered in vitro procedures in Markram et al. (1997b); Sjöström and Häusser (2006) and reproduced them in silico, as described in Section A.2. We then used a multi-objective genetic algorithm (GA) Van Geit et al. (2016) to find model parameters best matching mean EPSP ratio *in silico* vs *in vitro*. To minimize the computational cost, we considered only 5 stimulation protocols, as summarized in Table 2.

**Table 2:** Stimulation protocols used for model fitting.

| Connection type | Frequency (Hz) | $\Delta t$ (ms) | Reference |
| --- | --- | --- | --- |
| L5 TTPC – L5 TTPC | 2 | 5 | Markram et al. (1997b) |
| L5 TTPC – L5 TTPC | 5 | 5 | Markram et al. (1997b) |
| L5 TTPC – L5 TTPC | 10 | 10 | Markram et al. (1997b) |
| L5 TTPC – L5 TTPC | 10 | -10 | Markram et al. (1997b) |
| L23 PC – L5 TTPC | 50 | 10 | Sjöström and Häusser (2006) |

**Table 3:**
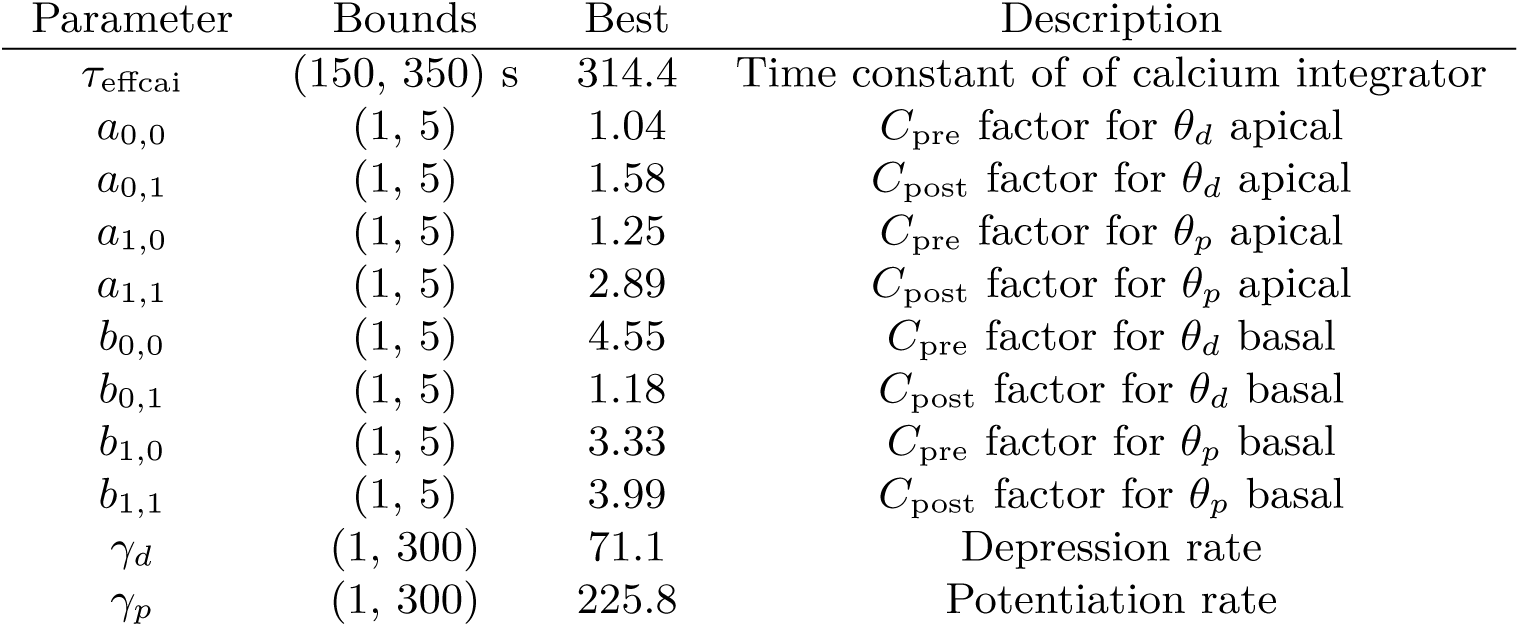
Optimized parameters with respective boundaries and best solution for the long term plasticity model.

The final solution found by the multi-objective genetic algorithm (MOGA) (see Table 3) was then validated by fully simulating all available protocols on a new set of connections, unseen in training.

## B. Supplementary results

**Figure B.1:**
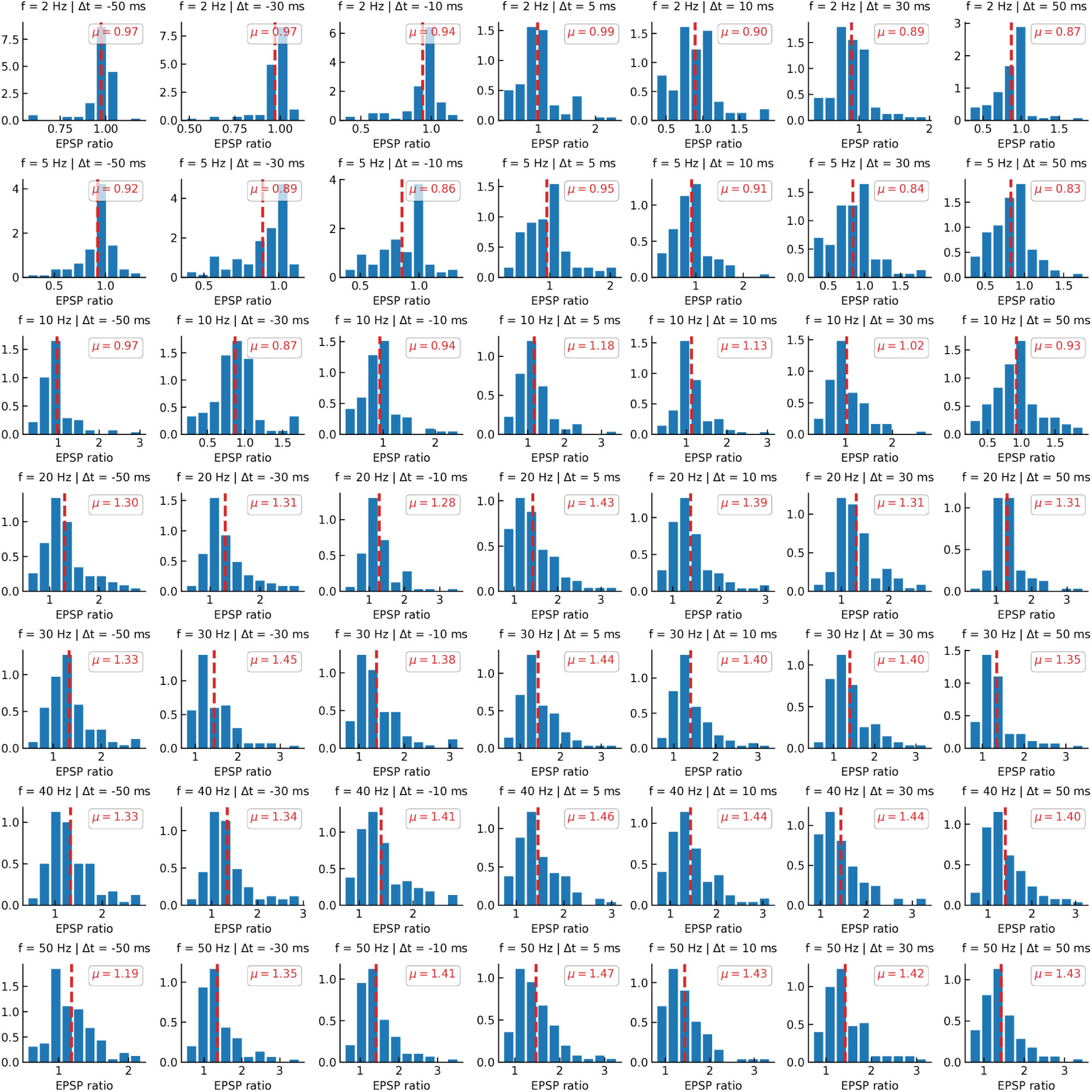
L5 PC – L5 PC plasticity experiment results.

**Figure B.2:**
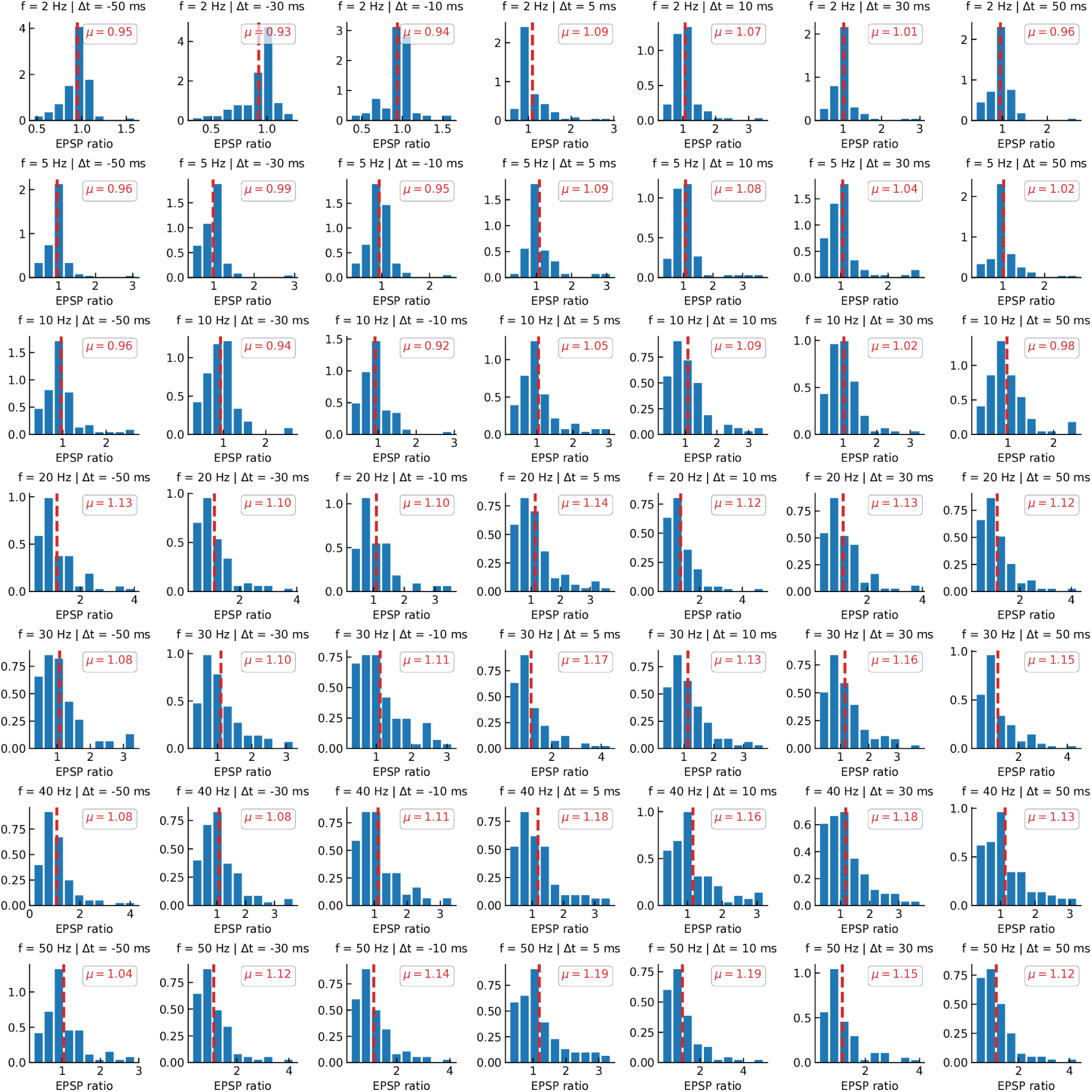
L23 PC – L5 PC plasticity experiment results.

**Figure B.3:**
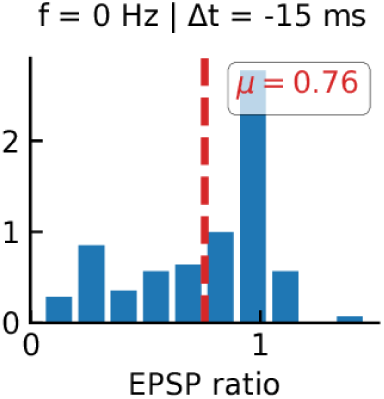
L4 PC – L23 PC plasticity experiment results.

**Figure B.4:**
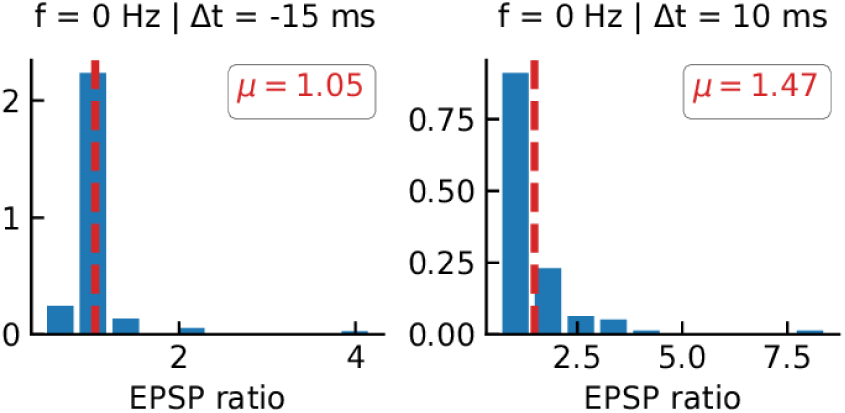
L4 PC – L23 PC plasticity experiment results, presynaptic MK801.

**Figure B.5:**
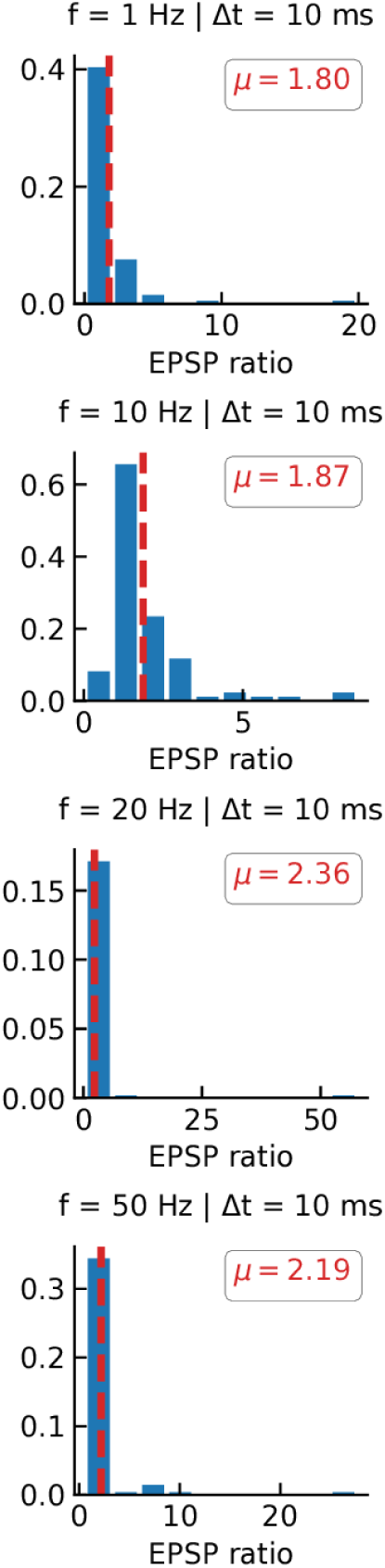
L23 PC – L23 PC plasticity experiment results.

**Figure B.6:**
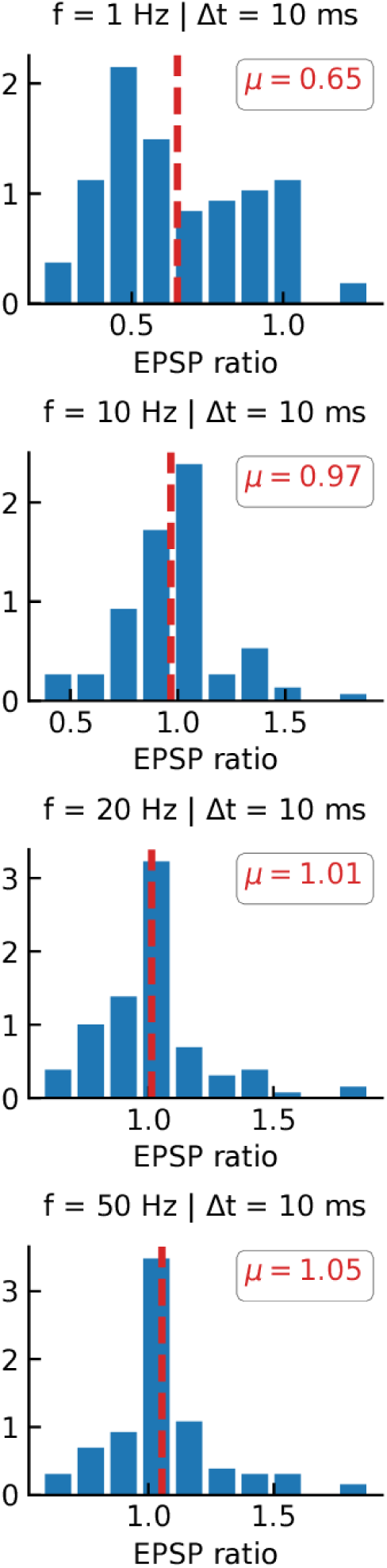
L4 SSC – L4 SSC plasticity experiment results.

**Figure B.7:**
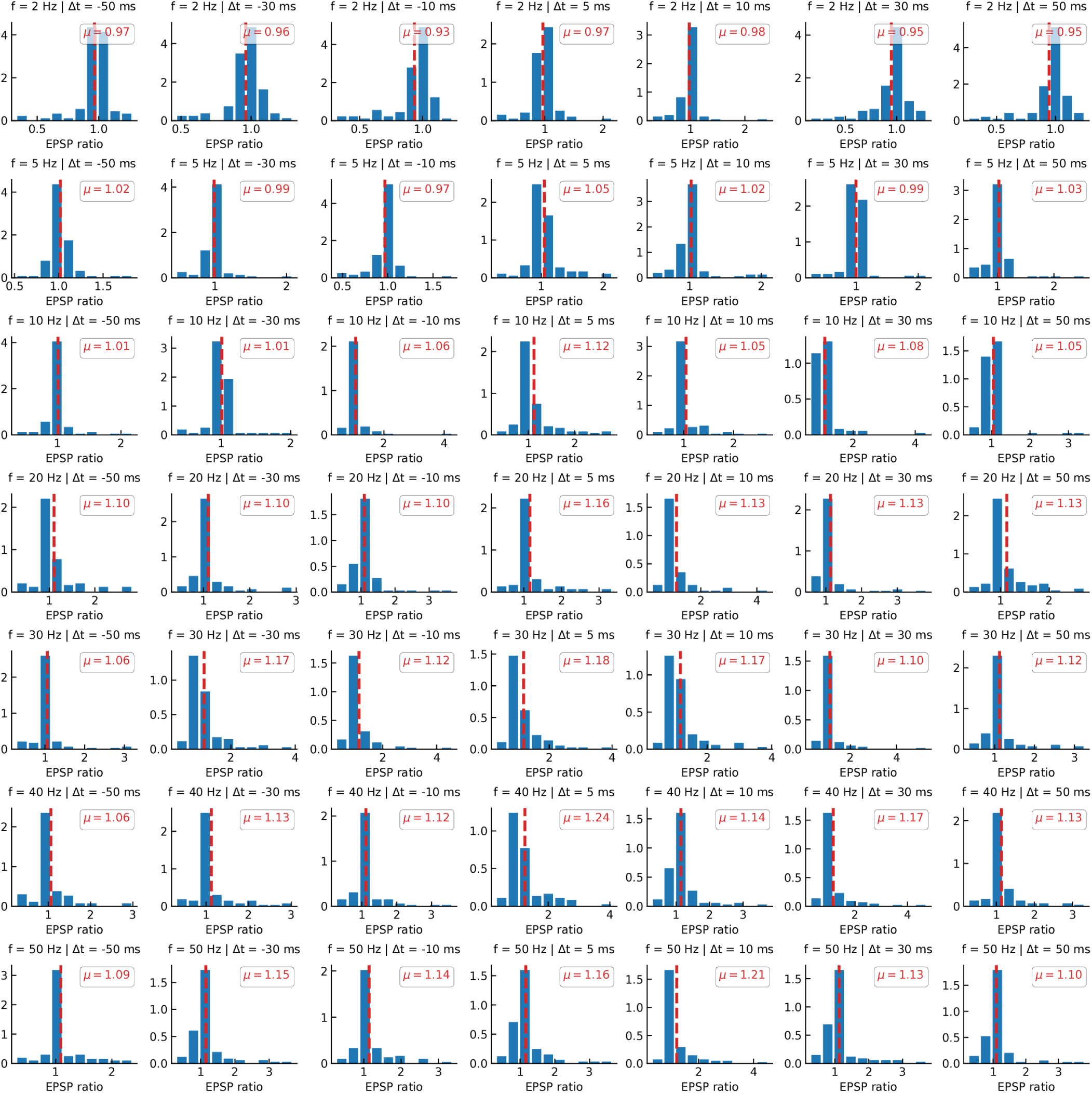
L4 PC – L6 PC plasticity experiment results.

**Figure B.8:**
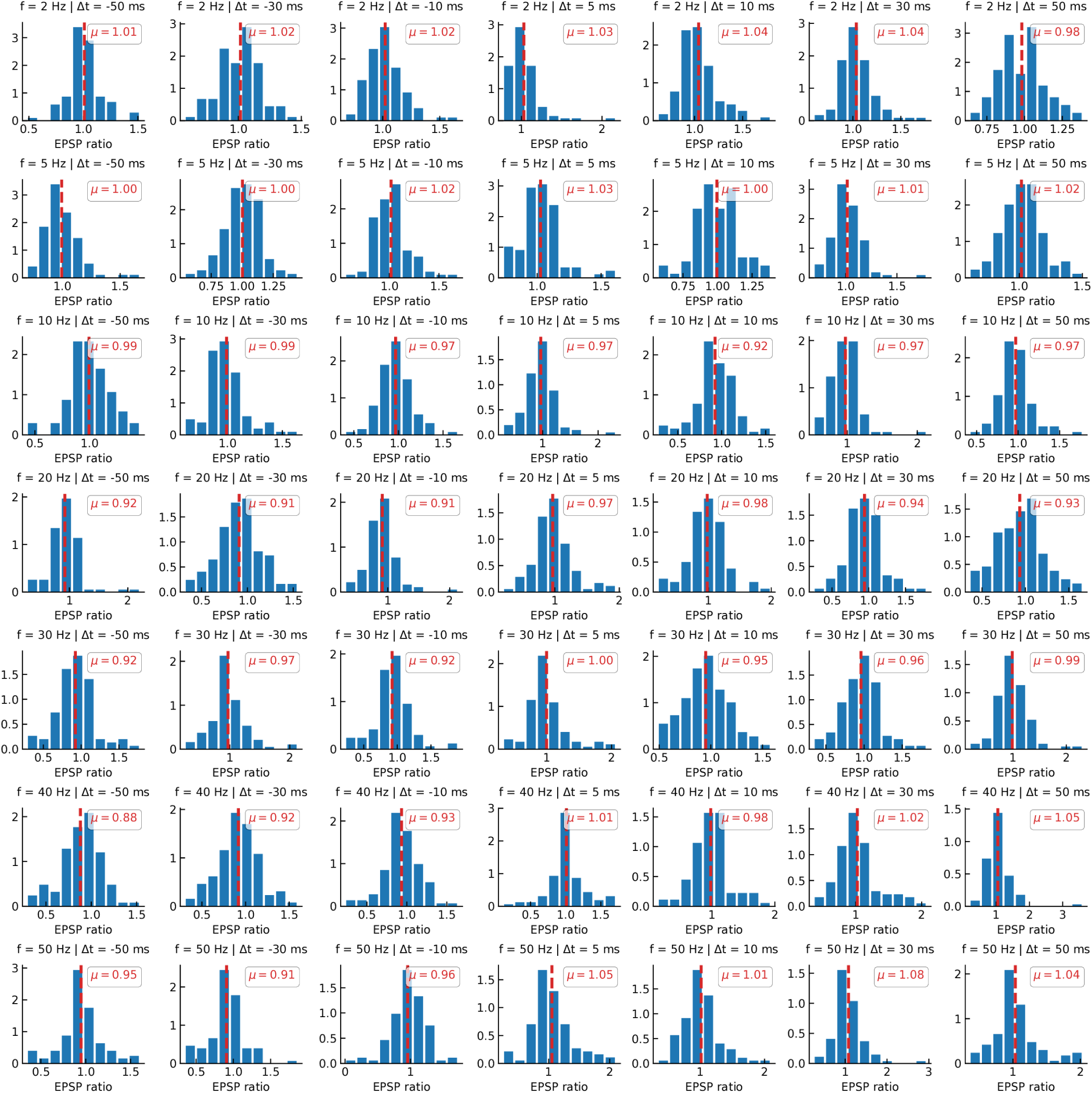
L5 PC – L5 PC plasticity experiment results, low calcium.

